# Deep learning regression model for antimicrobial peptide design

**DOI:** 10.1101/692681

**Authors:** Jacob Witten, Zack Witten

## Abstract

Antimicrobial peptides (AMPs) are naturally occurring or synthetic peptides that show promise for treating antibiotic-resistant pathogens. Machine learning techniques are increasingly used to identify naturally occurring AMPs, but there is a dearth of purely computational methods to design novel effective AMPs, which would speed AMP development. We collected a large database, Giant Repository of AMP Activities (GRAMPA), containing AMP sequences and associated MICs. We designed a convolutional neural network to perform combined classification and regression on peptide sequences to quantitatively predict AMP activity against *Escherichia coli*. Our predictions outperformed the state of the art at AMP classification and were also effective at regression, for which there were no publicly available comparisons. We then used our model to design novel AMPs and experimentally demonstrated activity of these AMPs against the pathogens *E. coli, Pseudomonas aeruginosa*, and *Staphylococcus aureus*. Data, code, and neural network architecture and parameters are available at https://github.com/zswitten/Antimicrobial-Peptides.

## 1 Introduction

Resistance to small molecule antibiotics is a growing public health concern. Antimicrobial peptides, or AMPs, are one strategy to address this issue. AMPs are short peptides that are a component of many animals’ innate immune systems. While they have multiple physiological functions (Hancock *et al.*, 2016), the best-studied function of AMPs is as broad-spectrum antibacterial agents (Mahlapuu *et al.*, 2016). One diverse group of AMPs, cationic AMPs or CAMPs, has been particularly well studied.

Since many CAMP sequences are known, and CAMPs mostly share biophysical properties and a membrane disruption-based mechanism of action (Nguyen *et al.*, 2011; Lee *et al.*, 2015), machine learning can be effective for CAMP discovery and design. For example, classification algorithms have been used to predict whether peptide sequences will be antimicrobial or not, which allows for scanning of sequenced genomes for antimicrobial peptide discovery. Such algorithms include AMP Scanner v2 (Veltri *et al.*, 2018), iAMPpred (Meher *et al.*, 2017), and a variety of algorithms available from the CAMP (Cationic AMP) database (Thomas *et al.*, 2009). Other groups have used regression approaches, based on peptide structure and biophysical properties, to quantitatively predict antimicrobial activity. These approaches are often used for local sequence optimization around a specific known AMP scaffold (Yoshida *et al.*, 2018; Hilpert *et al.*, 2006).

Beyond identifying and optimizing existing AMPs, several groups have used variational autoencoders (Das *et al.*, 2018) or generative recurrent neural network (RNN)-based models (Müller *et al.*, 2018; Nagarajan *et al.*, 2018) to generate new AMP sequences. These models generate sequences without an associated prediction of activity, although Nagarajan *et al.* further added a regression model (performance unspecified) to filter the designed sequences by predicted activity.

Our goal was to improve on these approaches by combining a large dataset with a regression model to design AMPs with a low predicted minimum inhibitory concentration (MIC). MIC is a standard measure of antibiotic activity: lower MIC means a lower drug concentration required to inhibit bacterial growth. We first assembled a large dataset of MIC measurements by combining data from multiple databases into GRAMPA (Giant Repository of AMP Activity).^1^ Examination of this dataset yielded experimental corroboration that MIC is more correlated among bacteria in the same gram class.

Next, we used GRAMPA to train convolutional neural network (CNN) models for AMP activity prediction. As a benchmark, we showed that converting our model to a classifier yields classification performance that improves on the state of the art.

Finally, we used simulated annealing over peptide space to design novel AMPs with low predicted MIC against *Escherichia coli*. Analysis of the model’s preferred sequences showed that it learned the concept of alpha-helical hydrophobic moment, a key signature of an active CAMP (Lee *et al.*, 2015). We designed two novel AMPs, and verified in vitro their potent activity against *E. coli* and also against *Pseudomonas aeruginosa* and *Staphylococcus aureus*, two important pathogens.

## 2 Methods

### 2.1 Data gathering and preprocessing

#### GRAMPA

We scraped all data from APD (Wang *et al.*, 2015), DADP (Novković *et al.*, 2012; this database appears to no longer be maintained), DBAASP (Pirtskhalava *et al.*, 2015), DRAMP (Fan *et al.*, 2016), and YADAMP (Piotto *et al.*, 2012). Each database was scraped in Spring 2018.

GRAMPA contains 6760 unique sequences, and 51345 total MIC measurements. Some peptide/bacteria pairs occurred multiple times due to overlap between databases and/or activity tested against multiple bacterial strains. To facilitate the reuse of our data without the need for the costly regex parsing and web scraping involved in amalgamating MIC data, we have made GRAMPA publically available on Github in the form of a single CSV with bacteria species and strain, AMP sequence and modification information, and a link to the original database entry.^1^

#### Preprocessing

We first excluded all sequences in GRAMPA with modifications other than standard c-terminal amidation and disulfide bonds. This meant excluding all data from YADAMP, which did not provide modification information. Where multiple measurements for a bacterium-AMP pair were present in our database, we took the geometric mean. Our preprocessed data contained 4559 peptides with associated log MIC values against *E. coli*, of which 3404 contain no cysteines and 1155 contain at least one cysteine. We split off 509 AMPs (15% of the no-cysteine dataset) for a held-out test set, and then removed the AMPs with a length >46 (which had sequences truncated) as we are not planning on designing AMPs of that length. This left 499 AMPs in the held-out test set. In order to select and tune a neural network architecture (Section 3.2), we split the remaining 2895 AMPs into a training set of size 2316 and a validation dataset of size 579. After selecting an algorithm, we recombined these data for a full training set of 2895 AMPs, or 4050 AMPs including those with cysteine.

#### Negative data from UniProt

In accordance with previous work (Veltri *et al.*, 2018), we generated a negative dataset from UniProt (The UniProt Consortium, 2018) by filtering for sequences with experimentally validated cytoplasmic localization and none of the terms “antimicrobial”, “antibiotic”, “antiviral”, “antifungal”, “secreted”, “excreted”, or “effector” (search string “locations:(location:”Cytoplasm [SL-0086]” evidence:experimental) NOT antimicrobial NOT antibiotic NOT antiviral NOT antifungal NOT secreted NOT excreted NOT effector”) We then filtered the sequences sharing >40% sequence identity using CD-HIT (Huang *et al.*, 2010) (the results of this filtering can be found at http://weizhong-lab.ucsd.edu/cdhit-web-server/cgi-bin/result.cgi?JOBID=1545509860). For each peptide in the positive dataset, we generated a negative peptide by selecting a random length-matched and cysteine-free substring from one of these filtered non-antimicrobial sequences. Thus, our final set of negative data had exactly the same length distribution as the positive data.

### 2.2 Machine learning model designs

#### Peptide encoding

Amino acids were represented using a one-hot encoding, meaning that each amino acid was a vector of length 21 (20 amino acids and a 21^st^ entry for c-terminal amidation) where every entry is 0 except for a 1 at the index of the amino acid of interest, and a 1 at the 21^st^ position if the peptide is c-terminal amidated. A peptide was then encoded as a 21×46 matrix, where 46 was the maximum peptide length we accepted, so chosen because it marks the 95^th^ percentile of peptide length. Peptides shorter than the maximum length were padded with vectors of 21 zeros each; peptides longer than the maximum length were truncated.

#### Regularized linear model

Ridge regression was performed using the RidgeCV module of the Python sklearn package (Pedregosa *et al.*, 2011), using leave-one-out cross-validation. Regularization parameter α was optimized to the nearest integer value. We trained two ridge regression models, each with a different peptide featurization. The first featurization was simply the amino acid compositions of the peptides, a vector of length 19 (since cysteine was not included), and the second was a flattened one-hot encoding vector consisting of 19×46=874 binary features.

#### *k*-NN analysis and sequence alignment

*k*-NN regression was performed defining “nearest neighbor” in one of five ways. The first was the edit, or Levenshtein, distance (Levenshtein, 1966). For the other four, we varied two factors in calculating the similarity between a query AMP and the AMPs in the training set: the alignment type (local or global) and the scoring matrix used (identity matrix vs. PAM30 substitution matrix). Alignment scores were calculated using the “pairwise2” function in the Biopython Python package (Cock *et al.*, 2009), the scoring matrices “matlist.ident” and “matlist.pam30”, a gap opening penalty of −9, and a gap extension penalty of −1. Local alignments using these gap penalties and the PAM30 matrix were also used to generate Figures 6 and S6.

For *k*-NN-based classification, we generated a length-matched random peptide negative training dataset. To predict the class of a query peptide, we had the *k* nearest neighbors (evaluated using Levenshtein distance as it was found to be superior to the other approaches in the regression analysis) “vote” based on whether they were AMPs or negatives. We had the best results with *k*=7.

#### Neural network model

For our architecture, after zero-padding, we begin with 2 1-dimensional convolutional layers of 64 neurons each with a kernel size 5 letters and a stride of 1 letter, paired with a Max Pooling layer with stride 2 and a pooling size of 2. We then use a flattening layer. Next, we add a Dropout(0.5) layer to regularize. Finally, we add two dense layers of 100 and 20 neurons each (with ReLU activation), and then a single neuron to transform the output into a single scalar value: the predicted log MIC value for the peptide. A diagram of our architecture is given in Figure 1. The model was trained to minimize mean squared error using the Adam optimizer. Different recurrent depths, kernel sizes, dropout rates, and learning rates were explored as described in Table S1; the CNN proved largely insensitive to these hyperparameters. We also explored replacing the convolutional layers with vanilla RNN layers, LSTM layers, and bidirectional recurrent LSTM layers.

**Figure 1.**
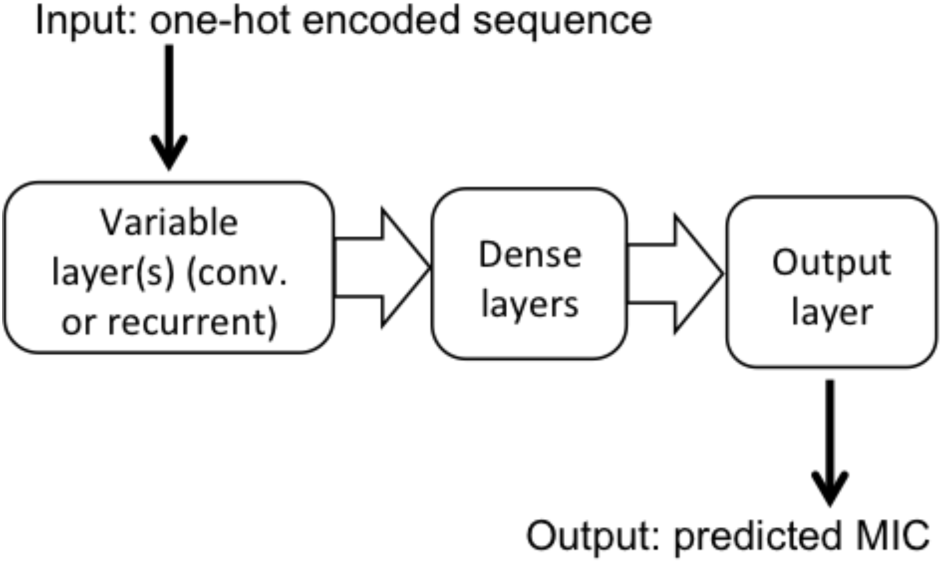
Architecture of neural network. Peptides are encoded as one-hot vectors and then fed to either convolutional or recurrent layers, followed by two dense layers and an output layer that outputs a predicted MIC for the peptide.

#### Negative Training Data

The majority of short amino acid sequences would have no antimicrobial activity if they were reified into real peptides. A model trained only on experimental data from existing peptide databases would have no inkling of this fact. We added negative training data to our model to reflect this prior, taking random sequences of amino acids and “labeling” them to have very low activity (log MIC = 4). We found doing so to increase the classification accuracy of the model, while slightly decreasing regression accuracy.

#### Ensemble model

To make our final model, we trained an ensemble of models with identical architecture on slightly different datasets. For positive datasts, we used the training set AMPs, and the training set AMPs with the cysteine-containing AMPs filtered out. We varied the amount of negative training data between 1, 3, and 10 times the size of the positive data, yielding a total of 2×3=6 different datasets. The random peptides in the negative data were allowed to include cysteine if and only if the positive dataset included cysteine. The networks were trained 5 different times for each negative dataset to average over the stochasticity inherent in training neural networks, meaning that the full ensemble model contained 6×5=30 neural networks.

We noticed while analyzing our model output that individual models gave extremely bimodal predictions, which was expected: the prediction was either very close to 4 (meaning, a predicted inactive peptide) or somewhere between −1 and 3.5 (meaning, a predicted active peptide). Therefore, for the purposes of classification (Section 3.3), instead of averaging over each of the ensemble model predictions, we had each model in the ensemble “vote.” If more than half of the models predicted log MIC > 3.9, we classified the peptide as inactive and predicted log MIC = 4. Otherwise, we classified the peptide as active and the predicted log MIC (used for generation of the ROC curves in place of a probabilistic prediction) was the average over all predictions that were <3.9. Finally, in our classification test sets we set all C-terminal amidation to “False” because the other algorithms did not have access to this information.

For regression and peptide design by simulated annealing, we simply averaged over the ensemble. Particularly for simulated annealing, it was important to have a smoother prediction landscape.

### 2.3 Simulated annealing for sequence design

Simulated annealing runs were initialized using a peptide with random sequence length between 10 and 25. Transitions were suggested according to the transition probabilities:

- 2.5% chance each of removing the residue at the beginning or end of the sequence
- 2.5% chance each of adding a random residue to the beginning or end of the sequence
- 90% chance of swapping a residue in the sequence (chosen randomly) for a randomly selected residue

The acceptance function was:

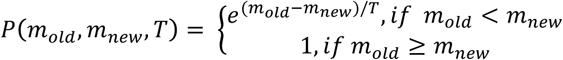

where *m*_*old*_ and *m*_*new*_ are the predicted log MIC values of the current peptide and new proposed peptide respectively, and *T* is temperature. A transition was also rejected if it did not satisfy the constraints we imposed, such as length (between 10 and 25) or charge density (see Section 3.5). The initial temperature *T*_*0*_ was set to 4/ln(2), meaning a transition probability of ½ for moving from an excellent peptide (log MIC = 0, or MIC = 1µM) peptide to an inactive peptide (log MIC = 4), and the final temperature *T*_*f*_ was 0.00001/ln(2). *N*_*steps*_ = 100,000 steps (transition proposals) were used per simulated annealing run, and temperature at step *n* varied according to the power law:

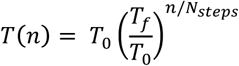

All generated peptides were a) cysteine-less (the random initializations and transitions only used the other 19 amino acids), and b) set to be c-terminal amidated, because they were later chemically synthesized with c-terminal amidation.

### 2.4 Hydrophobic moment analysis

Hydrophobic moments were calculated using the “Normalized consensus” residue hydrophobicities (Eisenberg *et al.*, 1984). Each peptide was compared to 1,000 randomly shuffled versions of itself to generate an “HM percentile,” defined as the frequency with which the peptide sequence has a greater HM than a randomly shuffled version of itself.

### 2.5 Minimum inhibitory concentration measurements

MIC values were measured using the broth microdilution method for cationic peptides (Wiegand *et al.*, 2008), with minor modifications. Peptides were synthesized at the Koch Institute Swanson Biotechnology Center. *P. aeruginosa* strain PAO1, *S. aureus* strain UAMS-1 and *E. coli* strain BL21 were grown overnight in BBL Mueller-Hinton II broth (MHB; BD Falcon) at 37°C. Peptide stock solution was prepared at 1mM in distilled water (CNN-SA1) and at 500 μM in distilled water with 0.1% acetic acid (CNN-SA2) then serially diluted 2-fold into 0.01% acetic acid, 0.2% Bovine Serum Albumin (BSA; Sigma Aldrich). The overnight cultures were diluted to a final concentration of approximately 3×10^4^ CFU/mL into fresh MHB. 90 μl of this inoculum was added to each well of a 96-well plate with 10 μl of the peptide dilution series, such that the final peptide concentrations evaluated were between 100 μM and 100 nM. After incubation at 37°C for 24 hours, the MIC of each peptide in MHB was determined by visually inspecting the plate to identify the lowest concentration at which there was no visible cellular growth. Reported values are the average of three technical replicates.

## 3 Results

### 3.1 Dataset characterization

Our dataset contained at least 700 MIC measurements for 10 different microbes, with the most measurements (4559) for *E. coli* (Table S2). To maximize our training set size, we selected *E. coli* as the organism against which we would train our model. Many AMPs were measured for their activity against multiple bacteria, which allowed us to consider the question of how tightly correlated the log MICs were between different microbial species. Since the primary mechanism of action of CAMPs is generally membrane disruption, and gram-negative bacteria have highly different membrane structures from gram-positive bacteria, we predicted that gram type would be the primary factor determining correlations. This trend was indeed observed in the data (Figure 2). We also included *Candida albicans*, an opportunistic pathogenic yeast, in our analysis. Antibiotic activity against *C. albicans*, a eukaryote, correlated poorly with antibiotic activity against all bacterial species (Figure 2). To our knowledge this is the first report on a large scale of AMP activity across multiple species. These results also confirm that an AMP designed for effectiveness against *E. coli* would also likely be effective against multiple gram-negative pathogens.

**Figure 2.**
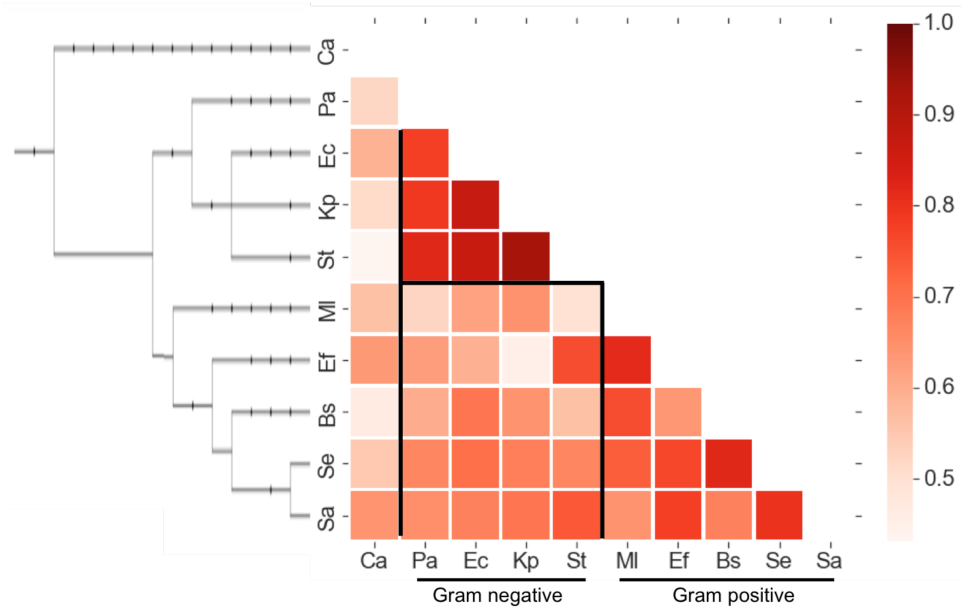
Pearson correlation of log MIC values for AMPs between different microbes. Left: phylogenetic tree, from PhyloT (http://phylot.biobyte.de/) and visualized using the interactive tree of life server (http://itol.embl.de/) (Letunic and Bork, 2016). Each correlation calculation included at least 50 MIC measurements. Black lines demarcate correlation blocks within a gram subtype, between gram types, and between *C. albicans* and a bacterial species. Abbreviations given in Table S2.

### 3.2 Design of machine learning architecture

After splitting the data into test and training sets, we considered multiple machine learning techniques and architectures. For the purposes of this section, we excluded data with cysteine and included no negative data for a simplified and streamlined comparison.

Before training the full ensemble, we optimized our model architecture by training a variety of networks with convolutional or recurrent layers. A performance comparison of the NN-based models on the validation set is given in Table S1. While many recent and state-of-the-art networks for AMP analysis are recurrent (Veltri *et al.*, 2018; Müller *et al.*, 2018), our convolutional neural network (CNN) model performed better than the recurrent models we tried. Additionally, model performance was not significantly altered by changing parameters such as dropout or convolutional kernel size (Table S1).

The superior performance of the CNN in comparison to recurrent architectures could be a reflection of the fact that many CAMPs are alpha helical in their active conformation (Lee *et al.*, 2015). Because alpha helices do not have long-range interactions, recurrent models may not be necessary for this problem. That said, it is possible that the recurrent models were simply slower to train and that with a larger dataset, network architectures that capture longer dependencies might start to show advantages.

We next compared our CNN’s performance to three baselines: two ridge regression (RR) models based on two different peptide featurizations (see Methods), and a k-nearest neighbors regression approach (Table 1). For the *k*-NN approach, we used the same training-validation split used to select a neural network architecture to select the best *k* and similarity measure (edit distance or alignment type and matrix; see Methods and Figure S1).

**Table 1.** Comparison of our CNN’s performance (no ensemble) with ridge regression and *k*-nearest neighbors. “Counts” denotes peptide featurization by amino acid counts only. “All” denotes one-hot encoding.

| Method | RMSE | Pearson $\rho$ | Kendall $\tau$ |
| --- | --- | --- | --- |
| CNN | 0.501 | 0.770 | 0.571 |
| Ridge (counts) | 0.644 | 0.560 | 0.379 |
| Ridge (all) | 0.602 | 0.627 | 0.432 |
| $k$-NN | 0.544 | 0.716 | 0.504 |

Our model was substantially better than both RR models, which suggested that it was successfully taking nonlinear and amino acid order effects into account and not simply basing predictions on amino acid composition. *k-*NN outperformed linear regression, but the deep learning model was the clear winner (Table 1).

We then trained the large ensemble model described in Methods for the results described below. This was because early investigation showed that while the ensemble was not much better than individual models for performance against held-out test sets, it was substantially better for sequence design. This was because our SA algorithm found spurious minima in the predicted log MIC landscape of single models: some sequences were predicted to be highly active by some models, but much less active by other models trained on the same data. The large ensemble model eliminated this issue and was thus used in all subsequent analysis.

### 3.3 Classification performance

While classification was not the primary purpose of this work, it allows for some benchmarking of our model compared to other machine learning-based AMP classifiers. We emphasize that the comparison is not specifically between the different machine learning algorithms so much as the classification capability of the data-algorithm combination. This is because the different models vary by training set size (1778 for AMP Scanner v2 (Veltri *et al.*, 2018), 2578 for the CAMP algorithms (Thomas *et al.*, 2009), 3417 for iAMPpred (Meher *et al.*, 2017), and 4050 for our model) and data labels (log MIC for our data, binary classification data for the other models).

We first evaluated classification performance of our CNN ensemble using random peptides as the negative data, and found that our classifier substantially outperformed other classifiers at this task, including AMP Scanner v2 which was previously shown to improve on the state of the art. This was also true for when we restricted the test set to peptides with no greater than 90% and 70% identity to the training set (Table 2 and Figure S2). We also tuned a *k-* NN classifier using the same training-validation split applied in Section 3.2 (Figure S3). As with regression, this classifier performed only slightly worse than our CNN (Table 2, Figure S2). It also performed better than all of the other classifiers except for CAMP-RF in the 70% identify filter case (Table S3a-c).

**Table 2.**
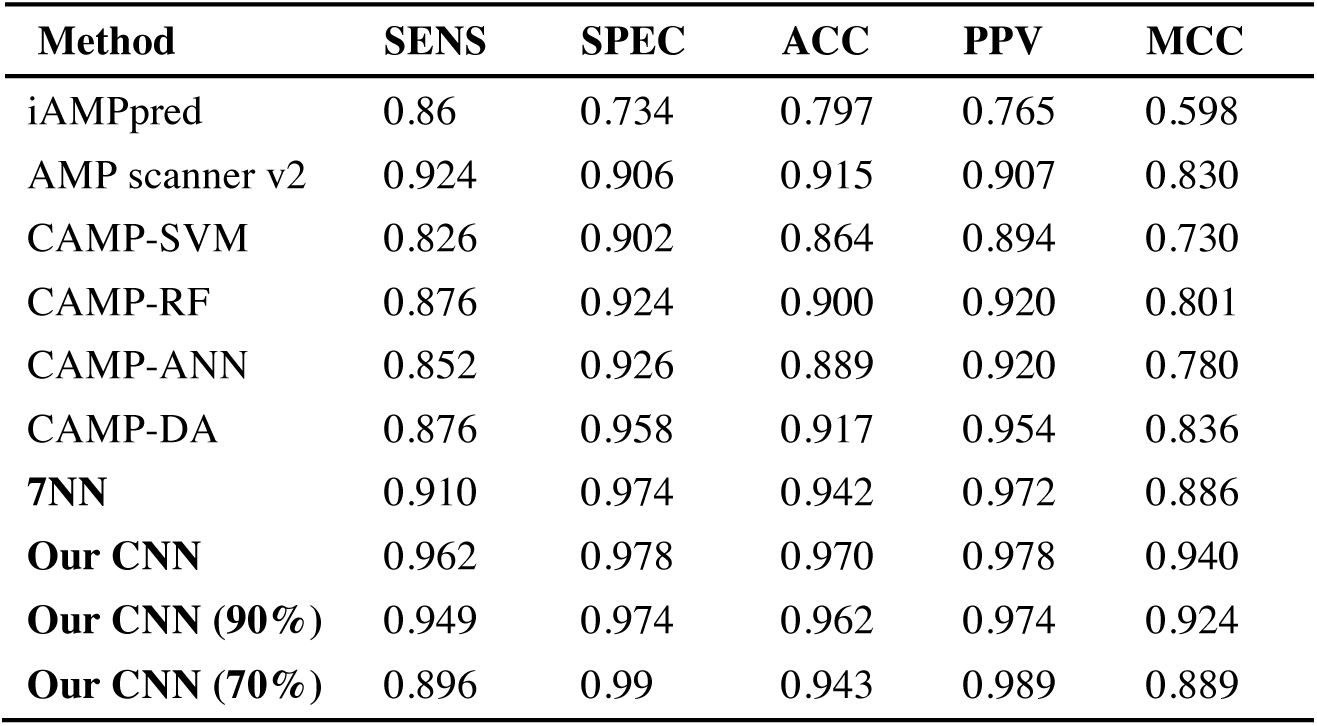
Classification performance comparison on AMP test set, using random peptides as negative data. SENS = sensitivity, SPEC = specificity, ACC = accuracy, PPV = positive predictive value (for test set of 50% positives, 50% negatives), MCC = Matthews Correlation Coefficient. 7NN = *k*-NN predictions (*k*=7). “Our CNN (90%)” and “Our CNN (70%)” rows show our CNN’s performance on modified test sets where AMPs sharing ≥ 90% or ≥ 70% sequence identity, respectively, with an AMP in the training set were removed.

| Method | SENS | SPEC | ACC | PPV | MCC |
| --- | --- | --- | --- | --- | --- |
| iAMPpred | 0.86 | 0.734 | 0.797 | 0.765 | 0.598 |
| AMP scanner v2 | 0.924 | 0.906 | 0.915 | 0.907 | 0.830 |
| CAMP-SVM | 0.826 | 0.902 | 0.864 | 0.894 | 0.730 |
| CAMP-RF | 0.876 | 0.924 | 0.900 | 0.920 | 0.801 |
| CAMP-ANN | 0.852 | 0.926 | 0.889 | 0.920 | 0.780 |
| CAMP-DA | 0.876 | 0.958 | 0.917 | 0.954 | 0.836 |
| **7NN** | 0.910 | 0.974 | 0.942 | 0.972 | 0.886 |
| **Our CNN** | 0.962 | 0.978 | 0.970 | 0.978 | 0.940 |
| **Our CNN (90%)** | 0.949 | 0.974 | 0.962 | 0.974 | 0.924 |
| **Our CNN (70%)** | 0.896 | 0.99 | 0.943 | 0.989 | 0.889 |

One important caveat is that other classifiers were trained using non-antimicrobial protein sequences from UniProt, as their negative data, not random peptides. This puts the other classifiers at a disadvantage for this comparison. UniProt sequences are more appropriate negative data if the goal is to scan genomes for antimicrobial sequences as protein sequences likely have different statistical properties from purely random peptides. While genome scanning was not our primary goal, we compared classification performance for our model and others against UniProt-derived negative data. This transfers the disadvantage to our model since we used random peptides, not UniProt sequences, as negative data. However, despite this handicap, our ensemble model had the second best performance of all of the tested models (after AMP Scanner v2) by Matthews Correlation Coefficient (MCC; Table S3 d-f) and area under the receiver operating characteristic curve (AUC; Figure S4). Notably, when we used a 10:1 negative:positive data ratio, the CNNs that emerged had the overall best classification performance, exceeding that of AMP Scanner v2 (Table S3 d-f). Nevertheless, we mixed these models with more balanced models to make the ensemble model we used for analyzing AMP candidates, as the ensemble demonstrated better regression performance.

The *k*-NN approach performed in the middle of the pack by AUC (Figure S4) and MCC except for particularly poor performance in the 70% identity case (Table S3d-f). While it is difficult to make concrete conclusions given the different goals and negative data of our model versus others, these comparisons along with *k*-NN’s good performance suggest that the improved net performance of our model derives mostly from our relatively large dataset. However, since *k*-NN performed particularly poorly on more unique AMPs, more complex predictive models will be better at designing novel, interesting sequences.

### 3.4 Regression performance

We next turned back to the problem of regression. Figure 3 depicts the predictions of our ensemble model on the AMP-only test set and Table S4 contains fit statistics. Our model’s predictions span over three orders of magnitude in MIC, which lent confidence that peptide design using this model will be likely to give particularly active AMPs. We do note that some active peptides were predicted to be inactive (log MIC ∼4) using our model, which suggests that at least some of AMP sequence space will be inaccessible to our design algorithm. This is due to the inclusion of negative data in our training set and accounts for the lower correlation coefficients observed here compared to the results in Table 1. Regardless, these false negatives are only a minor problem as we are not attempting an exhaustive search of all possible AMPs, just trying to find some that work well. More important is that there are very few peptides in the top left corner of the plots, meaning that the peptides predicted to be highly effective were, in fact, highly effective.

**Figure 3.**
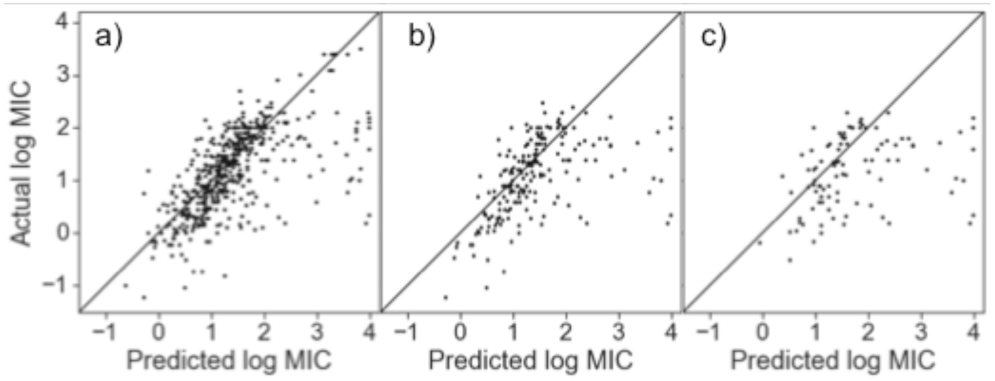
Predicted versus actual log MIC for peptides in test set with y=x line shown. AMPs sharing sequence identity ≥ some threshold with an AMP in the training set were removed: (a) no threshold, (b) 90%, (c) 70%.

### 3.5 Peptide design

We used simulated annealing to design peptide sequences with low predicted MIC values. Because high positive charge is believed to result in hemolysis and related toxicity towards eukaryotes and thus reduce the selectivity of AMPs, we imposed one of two different constraints on our sequence search to reduce the charge: a positive charge, and a positive charge density, constraint. In the first, we limited the total number of R’s and K’s to 6, and in the second, we permitted no more than 40% of the residues to be R or K. The generated peptides were predicted to have low MIC values, frequently below 1 μM (Figure S5).

To determine the extent to which our sequence design was capturing typical AMP structure, we analyzed the hydrophobic moment of the peptides assuming an alpha-helical conformation. The distribution of hydrophobic moments of our designed peptides did indeed show an elevated hydrophobic moment compared to shuffled versions of themselves (Figure 4). This elevated hydrophobic moment was not present when as a negative control we used a 140° turn per residue (Figure S6; as opposed to the 100° turn in alpha helices). Thus, our model learned to incorporate alpha helical structure into its sequence design.

**Figure 4.**
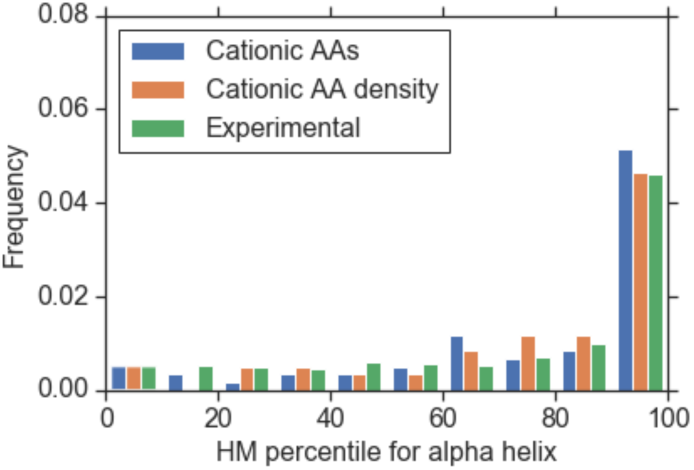
Histogram of “hydrophobic moment (HM) percentile” for designed peptides and experimental sequences from our dataset, using a 100° turn characteristic of alpha helices. HM percentile for a peptide is defined as the frequency with which the peptide sequence has a greater HM than a randomly shuffled version of itself.

Many sequences generated by simulated annealing constituted relatively minor variations on existing peptides. Figure S7 shows some representative sequence alignments of our peptides with peptides in the database, showing strong similarity in many cases. This similarity, and specifically the frequent repetitions of “LAK,” is likely due to the presence in the training data of several low-MIC peptides with LAK repeats.

Nevertheless, several peptides did represent new designs. By manually sorting through all of the peptides with predicted log MIC < 0, we identified two peptides with fairly low sequence similarity to any one peptide in particular, which we termed CNN-SA1 and CNN-SA2. CNN-SA1 was in some ways a modified fusion of two previously identified AMPs, while the second peptide was a mostly new design (Figure 5). Furthermore, each of these peptides had a high hydrophobic moment, as can be observed on helical wheel plots generated using HeliQuest (Gautier *et al.*, 2008) (Figure S8; HM percentiles 99.0 and 97.8 respectively), and predicted log MICs of −0.2 and −0.11 respectively. We selected these two peptides for solid-phase synthesis and experimental testing.

**Figure 5.**
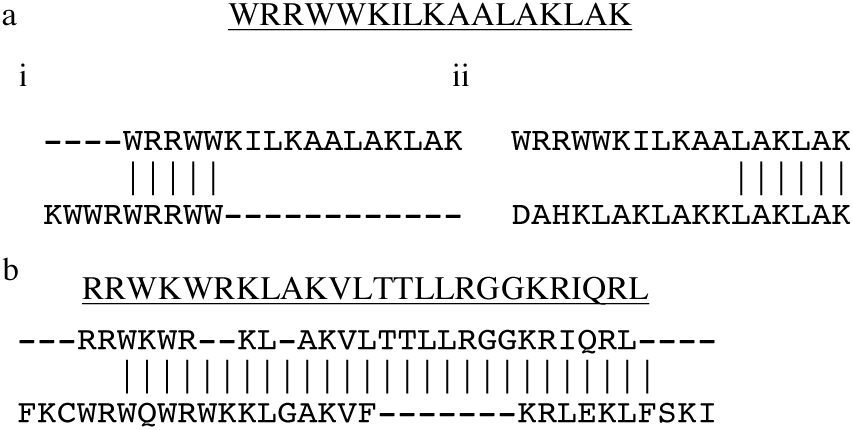
Designed peptides with limited similarity to peptides in the dataset, along with local alignments to peptides in dataset. a) CNN-SA1. i) Best local alignment to a peptide in dataset by PAM30 similarity matrix. ii) Representative alignment to a LAK-containing peptide. b) CNN-SA2 sequence with best alignment.

### 3.6 Experimental validation

Table 3 gives the antimicrobial activity of the two peptides. They have good activity against *E. coli*, as predicted: the MICs were lower than 79% of the active AMPs in our dataset. They also showed activity against *S. aureus* and *P. aeruginosa*, which is in line with what we would predict given that activities are usually correlated between different species of bacteria (Figure 2). Furthermore, we note that these peptides have not undergone experimental local sequence optimization, which frequently reduces MICs by an order of magnitude or more (López-Pérez et al., 2017). Thus, these sequences may be promising as lead compounds rather than as treatments in and of themselves.

**Table 3.** MICs (in μM) against various bacteria.

| AMP | *E. coli* | *P. aeruginosa* | *S. aureus* |
| --- | --- | --- | --- |
| CNN-SA1 | 3 | 8 | 25 |
| CNN-SA2 | 3 | 17 | 4 |

## 4 Discussion

We have developed a CNN-based model for predicting MIC values of antimicrobial peptides. While regression on activity has been performed before for very local variations about a single peptide, to our knowledge this is one of the first reports of global quantitative prediction of AMP activity and subsequent design of novel AMPs. When converted to a classifier, this CNN performed excellently at AMP recognition. Furthermore, we were able to use the CNN to design novel peptides with good predicted activity and hydrophobic moment. While most of the designed peptides showed strong similarity to already known AMPs, these peptides require no experimental effort to produce, so research attention can be concentrated on the ones that *are* unique. That said, future work will focus on ways to balance sequence diversity with predicted activity. Finally, another key design criterion that may be ripe for machine learning approaches is prediction of toxicity (Gautam *et al.*, 2014). If effective, computational toxicity prediction would be a critical step toward design of clinical useful AMPs.

## Acknowledgements

We thank Katharina Ribbeck and Rafael Gomez-Bombarelli for helpful discussions, and Tahoura Samad for helpful discussions and experimental help.

## Funding

J.W. was supported by the National Science Foundation Graduate Research Fellowship under Grant No. 1122374.

## Conflict of Interest

none declared.

## Footnotes

1 Available at https://github.com/zswitten/Antimicrobial-Peptides

