## Supplementary data for "Deep learning regression model for antimicrobial peptide design"

Jacob Witten,[1,*,†] Zack Witten[†]

[1]Department of Biological Engineering, Massachusetts Institute of Technology, Cambridge, MA 02139, USA.

[†] Co-first authors

**Table S1.** Comparison of different neural network architectures for MIC prediction. Layers: number of layers in the variable portion of the network, counting a convolutional-max pooling combination as one layer (the dropout and dense layers stay the same in each model). Kernel size: size of convolutional kernel (thus, n/a for recurrent networks). Dropout: level of dropout in the dropout layer. RMSE: root mean squared error for prediction of log MICs in validation set. Pearson $\rho$ and Kendall $\tau$: correlation coefficients between predicted and experimental log MICs in test set. Training and validation sets consist only of active AMPs without cysteine. CNNs performed best for regression. Model used for future analysis indicated in bold.

| Model | Layers | Kernel size | Dropout | Learning rate | RMSE | Pearson $\rho$ | Kendall $\tau$ |
|---|---|---|---|---|---|---|---|
| **CNN** | **2** | **5** | **0.5** | **0.001** | **0.544** | **0.723** | **0.516** |
| CNN | 2 | 1 | 0.5 | 0.001 | 0.595 | 0.666 | 0.472 |
| CNN | 2 | 2 | 0.5 | 0.001 | 0.550 | 0.710 | 0.514 |
| CNN | 2 | 3 | 0.5 | 0.001 | 0.567 | 0.691 | 0.482 |
| CNN | 2 | 4 | 0.5 | 0.001 | 0.553 | 0.708 | 0.497 |
| CNN | 2 | 6 | 0.5 | 0.001 | 0.550 | 0.718 | 0.505 |
| CNN | 2 | 7 | 0.5 | 0.001 | 0.562 | 0.697 | 0.490 |
| CNN | 1 | 5 | 0.5 | 0.001 | 0.562 | 0.701 | 0.503 |
| CNN | 2 | 5 | 0.9 | 0.001 | 0.557 | 0.707 | 0.517 |
| CNN | 2 | 5 | 0.7 | 0.001 | 0.550 | 0.713 | 0.508 |
| CNN | 2 | 5 | 0.3 | 0.001 | 0.563 | 0.696 | 0.492 |
| RNN | 1 | n/a | 0.5 | 0.001 | 0.628 | 0.648 | 0.456 |
| RNN | 2 | n/a | 0.5 | 0.001 | 0.713 | 0.529 | 0.355 |
| RNN | 3 | n/a | 0.5 | 0.001 | 0.657 | 0.599 | 0.406 |
| Bi-RNN | 1 | n/a | 0.5 | 0.001 | 0.651 | 0.615 | 0.423 |
| LSTM | 1 | n/a | 0.5 | 0.00001 | 0.662 | 0.531 | 0.354 |
| LSTM | 1 | n/a | 0.5 | 0.0001 | 0.660 | 0.591 | 0.408 |
| LSTM | 1 | n/a | 0.5 | 0.001 | 0.617 | 0.646 | 0.465 |
| LSTM | 1 | n/a | 0.5 | 0.01 | 0.608 | 0.666 | 0.464 |
| LSTM | 1 | n/a | 0.5 | 0.02 | 0.783 | 0.008 | 0.008 |
| Bi-LSTM | 1 | n/a | 0.5 | 0.001 | 0.610 | 0.657 | 0.459 |

**Table S2**. Number of AMPs with measured MIC by species (with abbreviations for Figure 2) following our preprocessing of GRAMPA as described in Section 2.1.

| Species | Sample size |
|---|---|
| *Escherichia coli* (Ec) | 4559 |
| *Staphylococcus aureus* (Sa) | 4110 |
| *Pseudomonas aeruginosa* (Pa) | 2516 |
| *Candida albicans* (Ca) | 1742 |
| *Bacillus subtilis* (Bs) | 1432 |
| *Staphylococcus epidermidis* (Se) | 947 |
| *Micrococcus luteus* (Ml) | 740 |
| *Klebsiella pneumoniae* (Kp) | 791 |
| *Salmonella typhimurium* (St) | 743 |
| *Enterococcus faecalis* (Ef) | 707 |
| All others | <310 |

**Table S3.** Classification performance on each of 6 different test sets. “7NN” used the kNN classification approach with our dataset and k=7. “No C” indicates our CNN model trained only on sequences without C; “w/C” indicates that sequences with C were allowed. “1x,” “3x,” and “10x” indicates the amount of negative data used to train that particular model, as a multiple of the amount of positive data (real AMP sequences). “All” indicates our full ensemble model incorporating all the sub-models. (a-c) Random peptides as negative test data. AMPs sharing sequence identity ≥ some threshold with an AMP in the training set were removed: (a) no threshold, (b) 90%, (c) 70%. (d-f) UniProt peptides as negative test data. AMPs sharing sequence identity ≥ some threshold with an AMP in the training set were removed: (d) no threshold, (e) 90%, (f) 70%. Best MCC in bold for each test set.

| **a)** | iAMP pred | AMP sc. v2 | CAMP -SVM | CAMP -RF | CAMP -ANN | CAMP -DA | 7NN | No C, 1x | No C, 3x | No C, 10x | w/C, 1x | w/C, 3x | w/C, 10x | All |
|---|---|---|---|---|---|---|---|---|---|---|---|---|---|---|
| SENS | 0.86 | 0.92 | 0.83 | 0.88 | 0.85 | 0.88 | 0.91 | 0.98 | 0.96 | 0.94 | 0.97 | 0.96 | 0.95 | 0.96 |
| SPEC | 0.73 | 0.91 | 0.90 | 0.92 | 0.93 | 0.96 | 0.97 | 0.92 | 0.97 | 0.99 | 0.92 | 0.98 | 0.99 | 0.98 |
| ACC | 0.80 | 0.91 | 0.86 | 0.90 | 0.89 | 0.92 | 0.94 | 0.95 | 0.97 | 0.97 | 0.95 | 0.97 | 0.97 | 0.97 |
| PPV | 0.76 | 0.91 | 0.89 | 0.92 | 0.92 | 0.95 | 0.97 | 0.92 | 0.97 | 0.99 | 0.93 | 0.98 | 0.99 | 0.98 |
| MCC | 0.60 | 0.83 | 0.73 | 0.80 | 0.78 | 0.84 | 0.89 | 0.90 | 0.93 | 0.93 | 0.90 | 0.94 | **0.94** | 0.94 |

| **b)** | iAMP pred | AMP sc. v2 | CAMP -SVM | CAMP -RF | CAMP -ANN | CAMP -DA | 7NN | No C, 1x | No C, 3x | No C, 10x | w/C, 1x | w/C, 3x | w/C, 10x | All |
|---|---|---|---|---|---|---|---|---|---|---|---|---|---|---|
| SENS | 0.86 | 0.93 | 0.86 | 0.91 | 0.87 | 0.89 | 0.90 | 0.96 | 0.94 | 0.93 | 0.96 | 0.95 | 0.93 | 0.95 |
| SPEC | 0.80 | 0.90 | 0.93 | 0.91 | 0.92 | 0.95 | 0.98 | 0.94 | 0.97 | 0.99 | 0.92 | 0.97 | 0.99 | 0.97 |
| ACC | 0.83 | 0.92 | 0.89 | 0.91 | 0.89 | 0.92 | 0.94 | 0.95 | 0.96 | 0.96 | 0.94 | 0.96 | 0.96 | 0.96 |
| PPV | 0.81 | 0.91 | 0.92 | 0.91 | 0.91 | 0.95 | 0.98 | 0.95 | 0.97 | 0.99 | 0.92 | 0.97 | 0.99 | 0.97 |
| MCC | 0.66 | 0.83 | 0.79 | 0.83 | 0.79 | 0.85 | 0.89 | 0.91 | 0.92 | 0.92 | 0.88 | 0.92 | **0.92** | 0.92 |

| **c)** | iAMP pred | AMP sc. v2 | CAMP -SVM | CAMP -RF | CAMP -ANN | CAMP -DA | 7NN | No C, 1x | No C, 3x | No C, 10x | w/C, 1x | w/C, 3x | w/C, 10x | All |
|---|---|---|---|---|---|---|---|---|---|---|---|---|---|---|
| SENS | 0.81 | 0.85 | 0.80 | 0.89 | 0.77 | 0.82 | 0.80 | 0.93 | 0.89 | 0.85 | 0.93 | 0.90 | 0.88 | 0.90 |
| SPEC | 0.89 | 0.89 | 0.94 | 0.95 | 0.95 | 0.92 | 0.99 | 0.95 | 0.99 | 0.99 | 0.97 | 0.98 | 0.99 | 0.99 |
| ACC | 0.85 | 0.87 | 0.87 | 0.92 | 0.86 | 0.87 | 0.90 | 0.94 | 0.94 | 0.92 | 0.95 | 0.94 | 0.93 | 0.94 |
| PPV | 0.88 | 0.88 | 0.93 | 0.94 | 0.94 | 0.91 | 0.99 | 0.95 | 0.99 | 0.99 | 0.97 | 0.98 | 0.99 | 0.99 |

| | | | | | | | | | | | | | | |
|---|---|---|---|---|---|---|---|---|---|---|---|---|---|---|
| MCC | 0.70 | 0.74 | 0.75 | 0.83 | 0.73 | 0.74 | 0.81 | 0.88 | 0.88 | 0.85 | **0.90** | 0.88 | 0.87 | 0.89 |

| **d)** | iAMP pred | AMP sc. v2 | CAMP -SVM | CAMP -RF | CAMP -ANN | CAMP -DA | 7NN | No C, 1x | No C, 3x | No C, 10x | w/C, 1x | w/C, 3x | w/C, 10x | All |
|---|---|---|---|---|---|---|---|---|---|---|---|---|---|---|
| SENS | 0.86 | 0.92 | 0.83 | 0.88 | 0.85 | 0.88 | 0.91 | 0.98 | 0.96 | 0.94 | 0.97 | 0.96 | 0.95 | 0.96 |
| SPEC | 0.89 | 0.93 | 0.87 | 0.93 | 0.85 | 0.90 | 0.89 | 0.66 | 0.87 | 0.96 | 0.65 | 0.84 | 0.94 | 0.86 |
| ACC | 0.87 | 0.93 | 0.85 | 0.90 | 0.85 | 0.89 | 0.90 | 0.82 | 0.92 | 0.95 | 0.81 | 0.90 | 0.94 | 0.91 |
| PPV | 0.88 | 0.93 | 0.86 | 0.92 | 0.85 | 0.90 | 0.89 | 0.74 | 0.88 | 0.96 | 0.74 | 0.85 | 0.94 | 0.87 |
| MCC | 0.75 | 0.85 | 0.70 | 0.80 | 0.71 | 0.78 | 0.80 | 0.68 | 0.84 | **0.89** | 0.66 | 0.80 | 0.89 | 0.83 |

| **e)** | iAMP pred | AMP sc. v2 | CAMP -SVM | CAMP -RF | CAMP -ANN | CAMP -DA | 7NN | No C, 1x | No C, 3x | No C, 10x | w/C, 1x | w/C, 3x | w/C, 10x | All |
|---|---|---|---|---|---|---|---|---|---|---|---|---|---|---|
| SENS | 0.86 | 0.93 | 0.86 | 0.91 | 0.87 | 0.89 | 0.90 | 0.96 | 0.94 | 0.93 | 0.96 | 0.95 | 0.93 | 0.95 |
| SPEC | 0.89 | 0.95 | 0.86 | 0.93 | 0.92 | 0.90 | 0.90 | 0.73 | 0.91 | 0.97 | 0.70 | 0.89 | 0.95 | 0.88 |
| ACC | 0.88 | 0.94 | 0.86 | 0.92 | 0.90 | 0.90 | 0.90 | 0.85 | 0.93 | 0.95 | 0.83 | 0.92 | 0.94 | 0.91 |
| PPV | 0.89 | 0.95 | 0.86 | 0.93 | 0.92 | 0.90 | 0.90 | 0.78 | 0.92 | 0.97 | 0.76 | 0.89 | 0.95 | 0.89 |
| MCC | 0.76 | 0.88 | 0.72 | 0.85 | 0.79 | 0.80 | 0.80 | 0.72 | 0.86 | **0.90** | 0.69 | 0.84 | 0.89 | 0.83 |

| **f)** | iAMP pred | AMP sc. v2 | CAMP -SVM | CAMP -RF | CAMP -ANN | CAMP -DA | 7NN | No C, 1x | No C, 3x | No C, 10x | w/C, 1x | w/C, 3x | w/C, 10x | All |
|---|---|---|---|---|---|---|---|---|---|---|---|---|---|---|
| SENS | 0.81 | 0.85 | 0.80 | 0.89 | 0.77 | 0.82 | 0.80 | 0.93 | 0.89 | 0.85 | 0.93 | 0.90 | 0.88 | 0.90 |
| SPEC | 0.91 | 0.96 | 0.89 | 0.92 | 0.91 | 0.92 | 0.85 | 0.70 | 0.88 | 0.93 | 0.70 | 0.85 | 0.92 | 0.84 |
| ACC | 0.86 | 0.91 | 0.84 | 0.90 | 0.84 | 0.87 | 0.83 | 0.81 | 0.88 | 0.89 | 0.81 | 0.88 | 0.90 | 0.87 |
| PPV | 0.90 | 0.95 | 0.88 | 0.91 | 0.89 | 0.91 | 0.85 | 0.75 | 0.88 | 0.92 | 0.75 | 0.86 | 0.91 | 0.85 |
| MCC | 0.72 | **0.82** | 0.69 | 0.80 | 0.68 | 0.74 | 0.66 | 0.64 | 0.76 | 0.78 | 0.64 | 0.75 | 0.79 | 0.74 |

**Table S4.** Performance of ensemble regression model on test sets unfiltered or filtered for identity with AMPs in the training set.

| Dataset | RMSE | Pearson $\rho$ | Kendall $\tau$ |
|---|---|---|---|
| All | 0.702 | 0.641 | 0.547 |
| 90% identity | 0.839 | 0.512 | 0.477 |
| 70% identity | 1.094 | 0.343 | 0.323 |

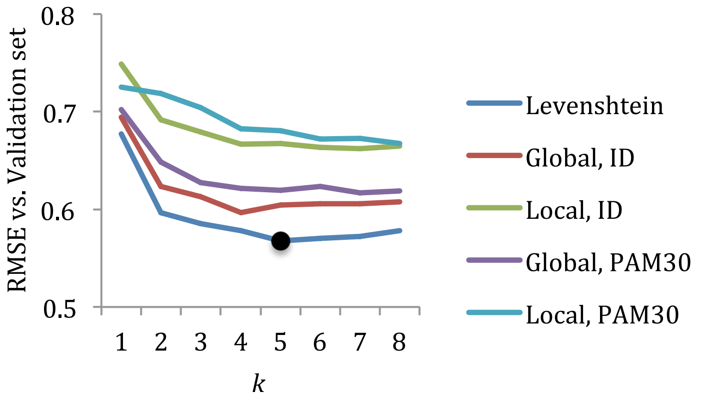


**Figure S1.** Root mean squared error (RMSE) of different *k*-nearest neighbors regression models on validation set. "Levenshtein" denotes standard Levenshtein distance, also known as edit distance, for identifying nearest neighbors. The other four lines use sequence alignment scores as similarity measures. "Global" and "local" denote type of sequence alignment used, "ID" or "PAM30" denote the similarity matrix used for the sequence alignment. Lowest RMSE indicated by black dot ($k = 5$, using edit distance for similarity).

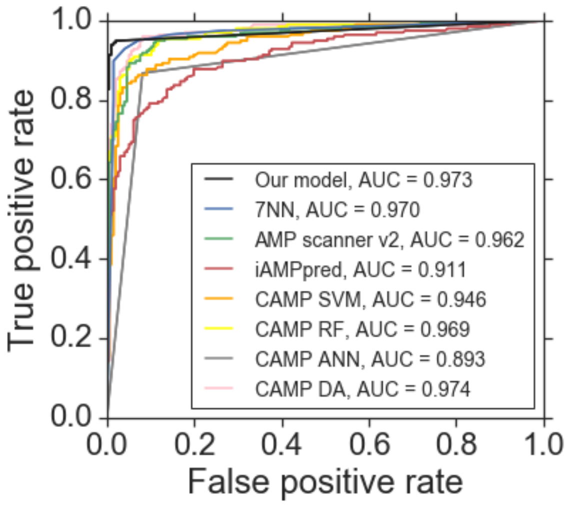


**Figure S2.** Receiver operating characteristic curve for classification performances of various predictive models on the test set, using **random** peptides as negative test data. AMPs sharing ≥ 90% sequence identity with an AMP in the training set were removed. AUC = area under receiving operator characteristic curve.

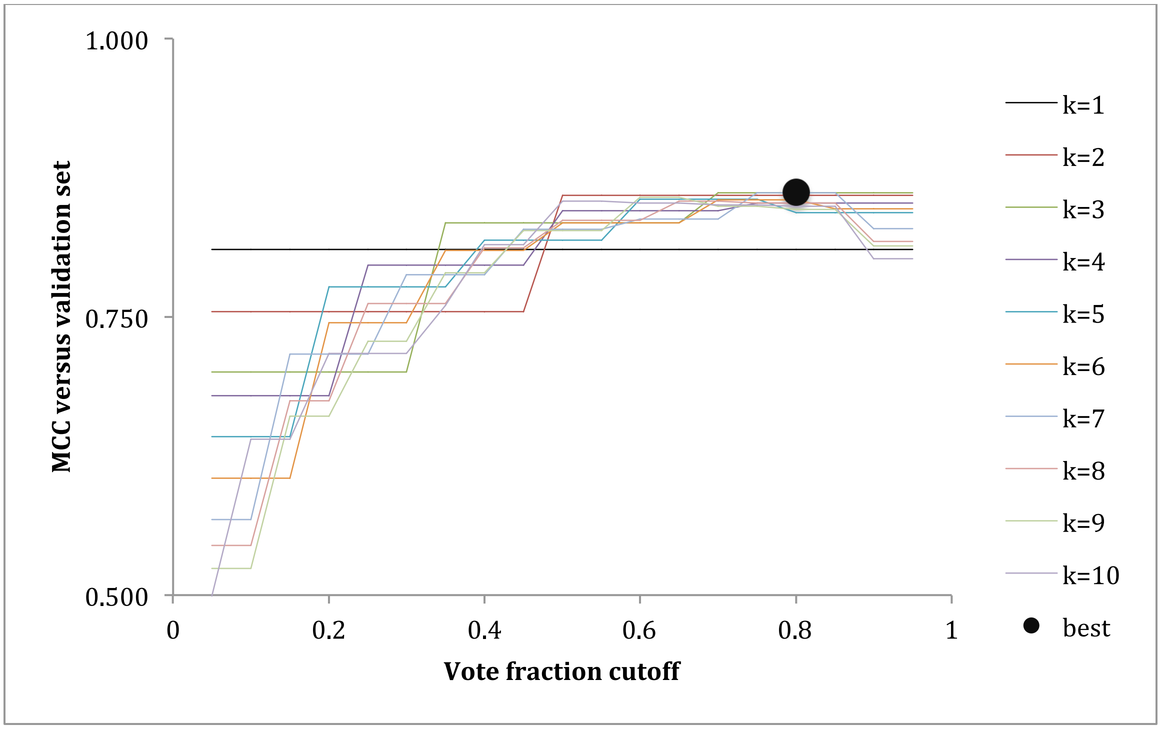


**Figure S3.** Hyperparameter optimization for *k*-NN classifier. Performance on validation set plus length-matched random peptides as negative data, as a function of *k* and the vote fraction cutoff *F*. A query peptide is predicted to be an AMP is the fraction $f_{query}$ of its *k* nearest neighbors that are AMPs is greater than *F*. Maximum MCC indicated by black dot (*k*=7, cutoff=0.8, MCC = 0.862); *k*=7 and 0.8 cutoff were used for future tests (as "7NN").

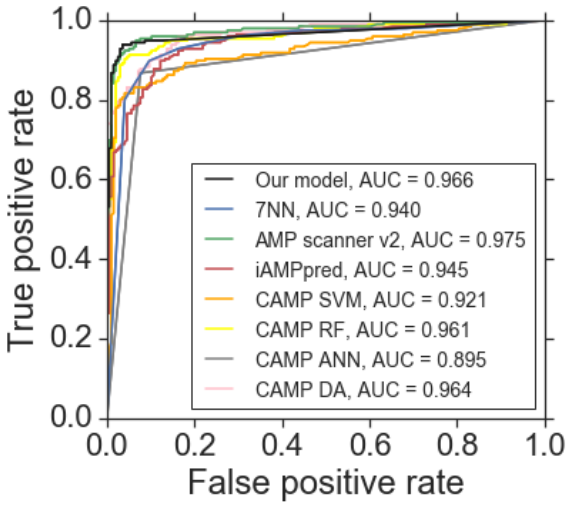


**Figure S4.** Receiver operating characteristic curve for classification performances of various predictive models on the test set, using **UniProt** peptides as negative test data. AMPs sharing ≥ 90% sequence identity with an AMP in the training set were removed. AUC = area under receiving operator characteristic curve.

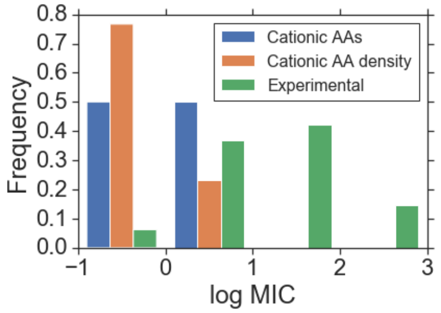


**Figure S5.** Histogram of predicted log MICs for designed peptides given a constraint on either the number (6; "Cationic AAs") or linear density (40%; "Cationic AA density") of cationic residues (R and K). Experimental log MICs from our dataset are given for comparison.

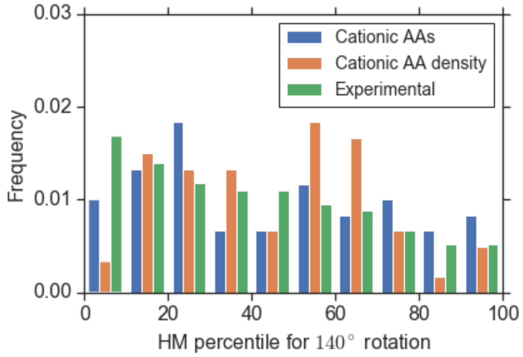


**Figure S6.** Histogram of "hydrophobic moment (HM) percentile" for designed peptides and experimental sequences from our dataset, using 140° turn per residue as a negative control. Percentiles are roughly uniformly distributed, unlike the bias toward high alpha-helical hydrophobic moment observed in Figure 5.

a
```
RKWIWWKLAKVLAKLAK
      |||||||||||
DAHKLAKLAKKLAKLAK
```

b
```
GFWKKILKKLATKLAKLAK
        |||||||||||
--DAHKLAKLAKKLAKLAK
```

c
```
GWKTLAKLAKKLAKLAK
    |||||||||||||
DAHKLAKLAKKLAKLAK
```

d
```
WRKFWKKILKKLAKLAKTGLRKPII---
    ||||||||||||||||||
---KWKKLLKKLLKLPKKLLKKLKKLLK
```

**Figure S7.** Example alignments of generated peptides (top of alignments) with peptides in the dataset (bottom of alignments), frequently showing the presence of "LAK" repeats and/or strong similarity to existing peptides.

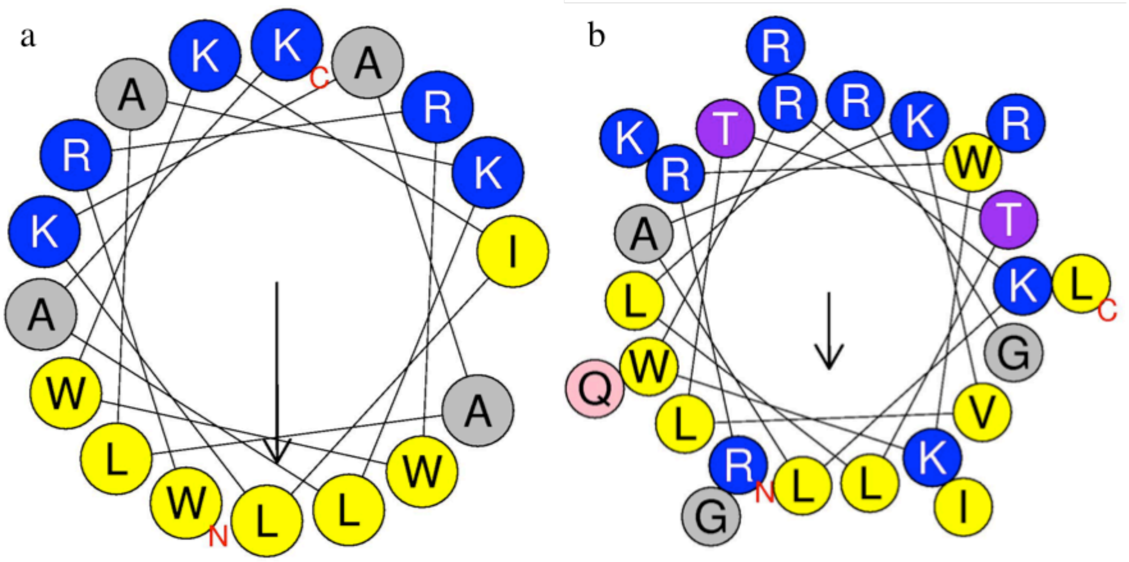


**Figure S8.** Helical wheel plot of two designed peptides a) CNN-SA1 and b) CNN-SA2, generated from HeliQuest (Gautier *et al*., 2008). Cationic residues in blue, hydrophobic residues in yellow. Both show pronounced alpha helical hydrophobic moment (represented by downwards arrow).
